# Fully unsupervised online spike sorting based on an artificial spiking neural network

**DOI:** 10.1101/236224

**Authors:** Marie Bernert, Blaise Yvert

## Abstract

Spike sorting is a crucial step of neural data processing widely used in neuroscience and neuroprosthetics. However, current methods remain not fully automatic and require heavy computations making them not embeddable in implantable devices. To overcome these limitations, we propose a novel method based on an artificial spiking neural network designed to process neural data online and completely automatically. An input layer continuously encodes the data stream into artificial spike trains, which are then processed by two further layers to output artificial trains of spikes reproducing the real spiking activity present in the input signal. The proposed method can be adapted to process several channels simultaneously in the case of tetrode recordings. It outperforms two existing algorithms at low SNR and has the advantage to be compatible with neuromorphic computing and the perspective of being embedded in very low-power analog systems for future implantable devices serving neurorehabilitation applications.

## Introduction

The collective dynamics of multiple single neurons constituting large neural ensembles underlies fundamental aspects of sensory^1,2^, motor^3–5^, navigation^6^, as well as cognitive functions^7^, and can be exploited for rehabilitation purposes^8^. Probing such dynamics relies on the identification and monitoring of the activity of a large number of individual cells at a high temporal resolution^9,10^. The activity of individual cells can be recorded over long periods of time (e.g., weeks or months) using extracellular microelectrodes and can be isolated using a spike sorting method^11^. Over the past decades, microfabrication technologies have made possible to build large-scale microelectrode arrays offering hundreds or thousands of recording sites in single probes^12–18^, and increasing efforts are currently put in the development of novel devices to make possible the recording of up to a million of cells simultaneously^19^. Yet, the amount of data generated by these very large-scale systems becomes a limiting factor to take advantage of all the information made available. A mandatory requirement is thus to find robust methods to process neural data online and without any user intervention (manual or semi-automatic intervention becoming impossible with a very high number of channels). These methods also need to be compatible with a low-power implementation in order to become eventually embeddable into the recording devices themselves. This is particularly crucial for implantable devices, which should satisfy the constraint to not heat the surrounding tissue, and also for wireless systems, for which the limited bandwidth precludes the transmission of whole raw data for very high numbers of channels. Ideally, intelligent implants would thus extract the relevant information already at the recording site before further transmission.

In this context, spike sorting remains a key issue. Although many different methods have been proposed^11, 20^, they still often require semi-automatic steps and high computation load not compatible with future low-power embedding. Most methods use three main separated steps to sort action potentials based on the discrimination between their different shapes originating from different neurons. The first step is action potential detection, usually done by a threshold-crossing criterion. This threshold can be applied directly on the signal or after suitable filtering, for example using an energy operator or templates^21^. The second step is feature extraction performed to reduce the dimensionality of the detected waveforms. This can be done using predefined features such as amplitude, width, wavelets coefficients^22^, or with dimensionality reduction algorithms such as PCA. The last and most computation-demanding step consists in separating the groups of features belonging to different action potentials using a clustering algorithm. Many clustering algorithms have been proposed, such as expectation-maximization on mixture of Gaussians^23^ or t-distributions^24^, K-means^25^, C-means^26^, superparamagnetic clustering^22^ or mean-shift^27^. Although some studies developed online spike-sorting methods^28,29^, most of the classical clustering algorithms require an offline processing step, as for instance online template matching^30^. Thus, fully automated and online spike-sorting remains a challenging problem. Moreover, the methods proposed so far require heavy computations not compatible with future analog embedding at the level of the recording site. In a first step toward low-power spike sorting, a neuromorphic architecture implementing a spike-timing-dependent plasticity (STDP) neural network has been developed for the clustering step^29^ but this network was not designed to fully process the continuous flow of incoming data by itself, requiring a separate detection stage.

Here, we propose a novel spike sorting method based on the use of an artificial spiking neural network implementing different plasticity rules. The key feature of this approach is to process continuously the stream of data recorded by a microelectrode and directly output trains of artificial spikes corresponding to the sorted activity of the recorded cells. By contrast with classical spike sorting approaches, this method does not use the conventional three-step procedure. Instead, the signal stream is continuously fed to the network, which rapidly learns to simultaneously detect and sort action potentials present in the signal thanks to its unsupervised learning properties. Once learning is stabilized, the output spike trains of the network predict those of the different active cells embedded in the input signal.

## Results

### Network architecture and behavior

We considered an artificial spiking neural network composed of three layers (input, intermediate and output) and one “attention” neuron, all connected by feedforward synapses implementing specific plasticity rules (Figure 1a). This network, composed of “sensory” and Leaky-Integrate-and-Fire (LIF) neurons, is bio-inspired rather than realistic. In particular the characteristic times are much shorter than realistic ones, which allows processing the information contained in the action potential shapes and makes the approach compatible with real-time spike sorting at the millisecond time scale. After learning, the output spikes of the network correspond to the detected and sorted action potentials present in the input signal. To avoid any confusion in the following, the term “spikes” thereafter denotes action potentials emitted by neurons of the artificial neural network, while “action potentials” denotes those of real neurons embedded in the input signal analyzed by the network.

**Figure 1.**
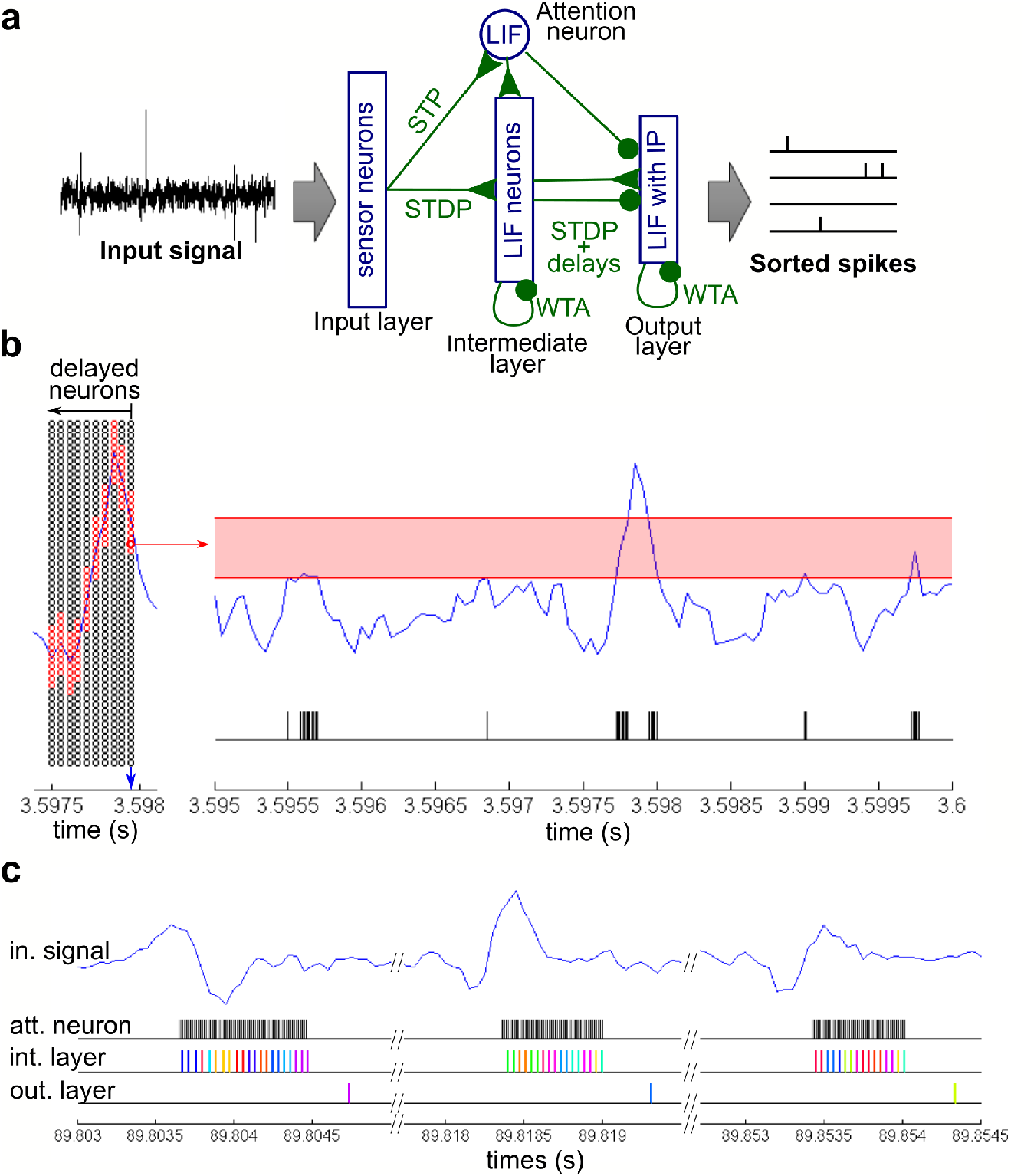
Network structure and behavior. **(a)** Overview of the network structure. Layers are connected in a feedforward manner. Each layer implements specific mechanisms and plasticity rules. Arrows with triangular endings represent excitatory synapses, while arrows with round endings represent inhibitory synapses. **(b)** The input layer encodes the incoming neural signal. Left panel: input layer activation at a specific time step (blue arrow), the red circles representing the activated input neurons and the black circles the inactive ones. Right panel: spike train emitted by one specific input neuron (highlighted in bold red on the left panel) for the input signal shown in blue. The red rectangle shows the sensitivity range of this input neuron, which fires when the input signal falls within this sensitivity range. **(c)** Example of the response of the different parts of the network for the occurrence of 3 different action potentials in the input signal (upper blue curve). The spike trains of the attention neuron, and of the intermediate and output layer neurons are shown below each input waveform. The different colors stand for different neurons within the same layer.

#### Input layer

A first layer, called the input layer, encodes the incoming neural signal within a sliding time window preceding the current time point into artificial spikes that are passed on to the next layers. Neurons of this input layer act as “sensory” neurons. They are organized as a grid where each column processes the signal at a different delay from the current time and, within each column, neurons are sensitive to different signal values and are set to fire at regular time steps when the input signal falls into its sensitivity range (Figure 1b). As a result, the input layer converts continuously the input signal into a set of artificial spike trains encoding at each time step the shape of the signal within a sliding window (Figure 1b and Supplementary Figure S1). The role of the rest of the network is to detect different patterns into this artificial input signal that reflect both the occurrences and the shapes of action potentials stemming from different cells.

#### Attention neuron

The role of the “attention” neuron is to detect every action potential occurrences embedded in the input signal. This neuron receives all spikes emitted by the input layer through excitatory synapses that implement a short-term plasticity (STP) rule. The STP rule weakens the weights of synapses for which presynaptic spikes occur at high frequency^31^. As a result, the weights of the synapses corresponding to presynaptic input neurons encoding signal amplitudes within the range of noise (and thus often activated) are weakened, while synapses corresponding to presynaptic input neurons encoding signal amplitude outside this range remain strong (see Supplementary Figure S2). These STP synapses thus make the “attention” neuron sensitive only to infrequent signal amplitudes and thus not to fire in response to noise but only when an action potential emerging from the noise occurs in the input signal (Figure 1c). The activity of the “attention” neuron in turn modulates the intermediate and output layers, thus acting as an attention mechanism gating the learning mechanisms occurring on these two subsequent layers.

#### Intermediate layer

The role of each intermediate layer neuron is to learn to recognize specific fragments of action potential waveforms present in the input signal so that the layer emits different firing sequences for different action potential waveforms (Figure 1c). The activity of intermediate layer neurons is gated by the attention neuron through fixed-weight excitatory synapses, ensuring that these neurons fire when the attention neuron fires and remain silent otherwise (Figure 1c). Neurons of the intermediate layer receive all spikes emitted by the input layer through excitatory synapses implementing an STDP rule (see Figure 2a and methods). The weights of these synapses are initialized randomly. Each time an intermediate neuron fires, the STDP rule is applied on its incoming synapses: the synapses stemming from the input neurons that were active during a given time window are potentiated while the others are depreciated. The STDP time window being short (equal to the delay between two input layer columns), the potentiated synapses reflect the shape of the signal at this specific time. Moreover, following a winner-take-all (WTA) mechanism, the active neuron also inhibits the other intermediate neurons so that it is the only one to fire and to reinforce its weights to learn this input layer pattern. Consequently, when a similar waveform fragment occurs again, this neuron is more likely to fire and thus reinforce the specificity of its response to this particular waveform pattern (Figure 2b&c). Once learning has been achieved, several intermediate neurons become specifically responsive to different waveform fragments (Figure 2d). The occurrence of an action potential in the input signal triggers a sequence of activation of intermediate neurons, which is different for different action potential waveforms and stable across occurrences of the same waveform (Figure 2e). These different sequences are then processed by the output layer, to sort the action potentials.

**Figure 2.**
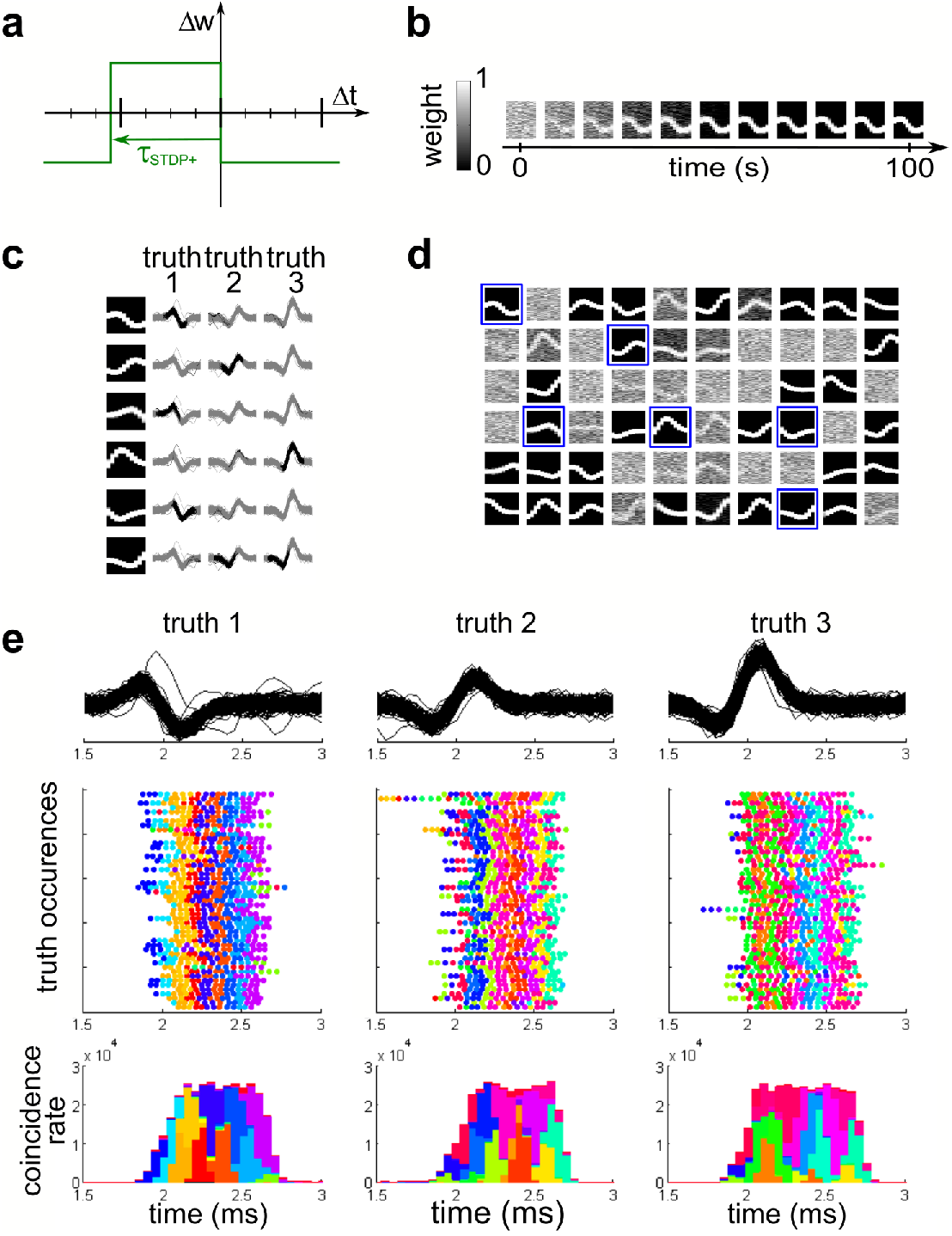
The intermediate layer learns waveform fragments. **(a)** STDP rule applied on the synapses stemming from the input layer. The weight change Δw depends on the time difference between the postsynaptic and presynaptic spike: Δt=t_pre_-t_post_. This rule is applied only when a postsynaptic spike occurs, so that no change happens if a presynaptic spike occurs alone. The main graduation shows the time interval between two input layer columns, and the secondary graduation shows the time interval between two input layer activations. **(b)** Weight evolution of the synapses projecting on one specific intermediate neuron. **(c)** Learnt shapes correspond to fragment of waveforms occurring in the signal. The first column shows the learnt weights for 6 different intermediate neurons (shown in d). The three other columns show the signal shape at each action potential occurrence, the black lines show the part of the signal encoded by the input layer at the moment when the intermediate neuron fires. **(d)** Weights learnt after a 200-s simulation by all neurons of the intermediate layer for an artificial dataset housing the 3 different action potential waveforms (see methods). Some neurons have learnt fragments of waveforms. The neurons squared in blue are the one shown in c, the first one is the one shown in **b. (e)** Timing consistency of the intermediate neurons’ spikes relative to the occurrences of each of the 3 action potential waveforms. Top row: signal shapes for all action potential occurrences on the last 100 s of simulation. Middle row: spike trains of the intermediate layer synchronized with 50 different action potential occurrences. Bottom row: distribution of intermediate spike times relative to each action potential occurrence (the histograms are cumulated). The different colors stand for different intermediate neurons.

#### Output layer

The purpose of the output layer is to finalize the spike-sorting process, by learning to recognize spike sequences produced by the intermediate layer that correspond to combinations of waveforms fragments. If the input signal contains action potentials of N different cells, a successful spike sorting results in exactly N neurons of the output layer each firing once for each occurrence of an action potential of a given cell in the input signal (Figure 1c). An important problem to overcome is the ability to learn patterns of different sizes and to differentiate between patterns that may overlap or include one into another. Overlapping pattern can occur for example if two action potential waveforms have the same beginning but different endings. The output layer was designed here to overcome these problems using four features in addition to the STDP rule (Figure 3a) and the WTA mechanism. First, the feedforward synapses are duplicated with different transmission delays, which give information to the output layer about the spike times of the intermediate layer (see Supplementary figure S3). Second, the output neurons are inhibited by the attention neuron, which forces them to wait until the end of a pattern before firing and thus to take into account the entire pattern to adjust the weights of incoming synapses (Figure 1c). Third, each neuron of the output layer receives the spikes of each neuron of the intermediate layer through both excitatory and inhibitory synapses implementing an STDP rule, allowing the overall weight to take negative or positive values. As a result, a neuron that has learnt a pattern gets excited by spikes belonging to this pattern and inhibited by spikes not belonging to the pattern. Thus, its potential is maximal when the pattern is exactly reproduced, without missing or additional intermediate neuron spikes (see supplementary figure S4). Fourth, the output neurons implement an intrinsic plasticity (IP) rule (Figure 3b), which enables them to adapt their threshold to the learnt pattern size. This ensures that an output neuron that has learnt to recognize a long pattern has a sufficiently high threshold preventing it to fire if the incoming pattern is incomplete, whereas a neuron that has learnt a short pattern has a low threshold and is still able to detect it. The introduction of this IP rule on the output neurons in conjunction with the use of excitatory and inhibitory synapses is key to solve the problem of recognition of overlapping patterns or patterns included one into another. The output neurons thresholds are initialized at a high value, and the feedforward synapses weights are initialized randomly with an average positive weight. Before learning, when an output neuron receives a spike pattern from the intermediate layer, it fires late due to its high threshold. According to the intrinsic plasticity rule, the threshold of a neuron learning a pattern progressively decreases until reaching an equilibrium value (Figure 3c), which can be shown to be proportional to the mean number of spikes received within the IP time window. As a consequence, the learning neuron fires earlier than the neurons that have not learnt any pattern. The synaptic weights evolve in parallel to the threshold (Figure 3c) according to the STDP rule. Once an output neuron has learnt a pattern (see neurons 3, 7 and 9 in the example of Figure 3d), most of its incoming synapses have converged to the minimum negative weight value. Only the synapses corresponding to the intermediate neurons and delays constituting the pattern converge to the maximum positive weight, and the few remaining synapses not relevant for discriminating the pattern converge a null weight (Figure 3d). By contrast, the incoming synaptic weights of neurons that did not learn any pattern remain distributed near the zero value. Overall, the proposed architecture of the network allows the output neurons to learn to recognize different patterns generated on the intermediate layer and to emit one spike for each occurrence of an action potential waveform in the input neural signal. Thus, each active output neuron directly predicts the activity of one real cell (Figure 4).

**Figure 3.**
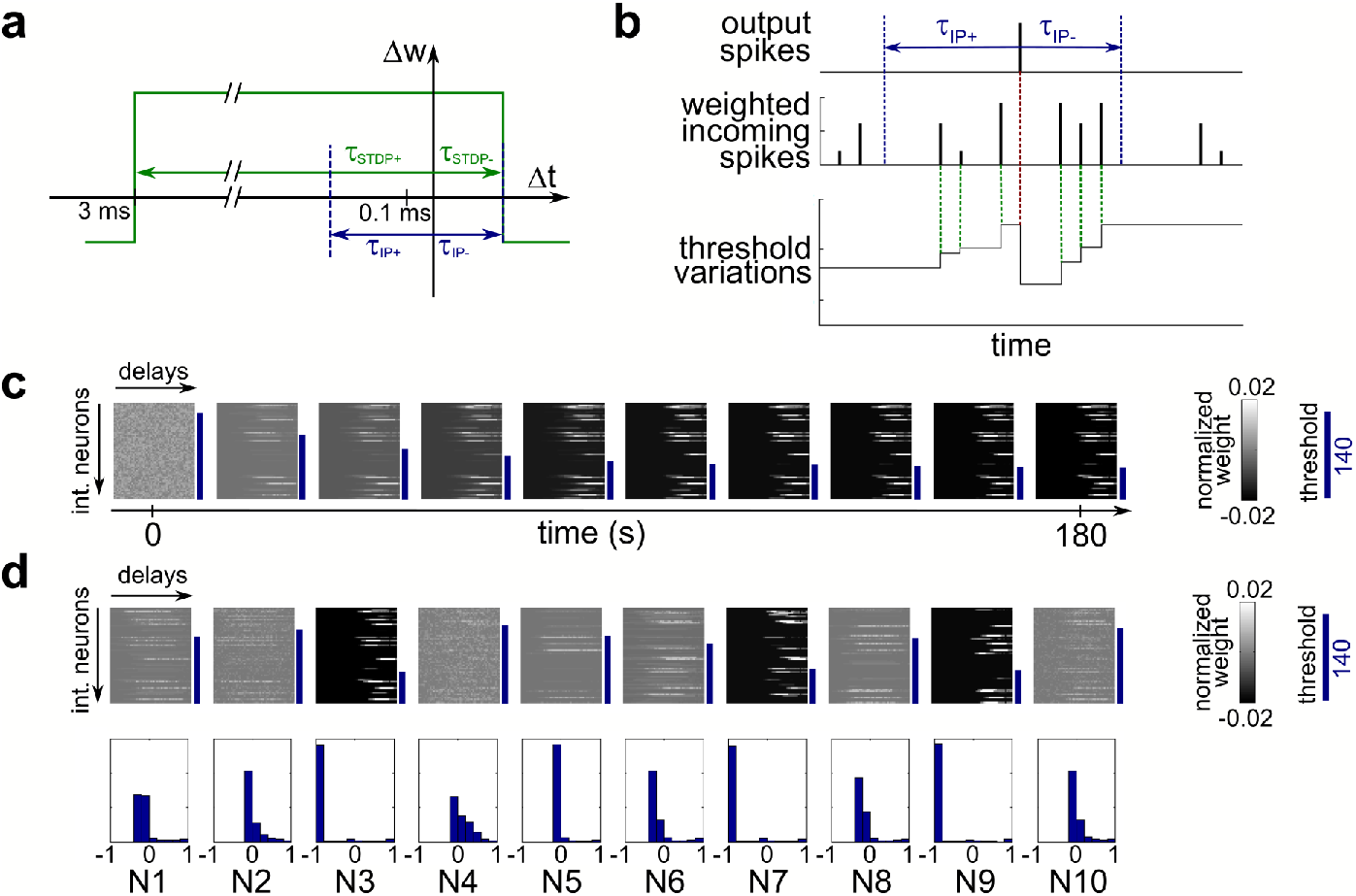
The output layer learns to recognize different intermediate layer patterns. **(a)** STDP rule applied on the synapses stemming from the intermediate layer. Similarly to the intermediate layer, the rule is applied for each postsynaptic spike occurrence, which means that no change happens if a presynaptic spike occurs alone. For the inhibitory synapses the sign of the weight change is reversed. **(b)** Intrinsic plasticity principle. Every time an output neuron receives a spike within a coincidence window around the postsynaptic spike, its threshold is increased proportionally to the weight of the synapse that transmitted the spike. Each time the neuron fires, the threshold is decreased proportionally to its own value. **(c)** Evolution of weights and thresholds. The matrices show the weight evolution of the synapses projecting on one output neuron. The synapses are organized according to their transmission delays and which neuron they stem from. Displayed normalized weights are the excitatory synapse weights minus the inhibitory synapse weights, divided by the neuron’s threshold. The blue bars shows the threshold evolution. **(d)** Synaptic weights learned by each of the 10 output neurons. Top row: final synaptic weights and thresholds at the end of a 200-s simulation, displayed similarly as in c. Bottom row: distributions of final synaptic weights (computed as the sum of the excitatory weight and the inhibitory weight) for each output neuron.

**Figure 4.**
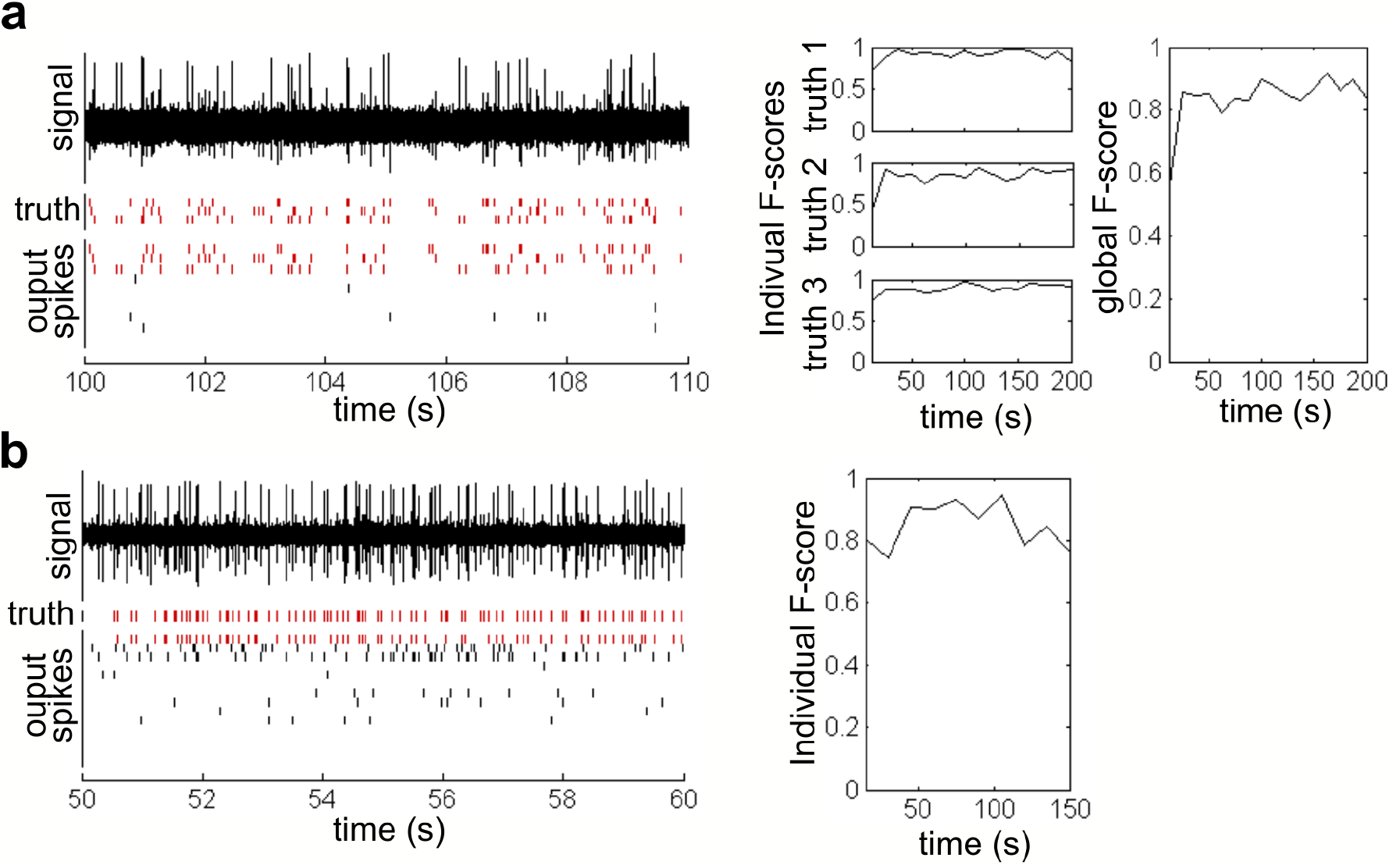
Examples of spike sorting achieved by the artificial network on simulated and real data. **(a)** Simulated data. Left: a 10-s segment of input signal is shown above the true spike trains for the 3 different waveforms and of the output spike trains of the artificial network. Output neurons have been reordered to visualize the similarity between the output spikes and the truth. Right: evolution of the performance of the network over the 200-s simulation. The F-score is computed both for the overall truth and output, and for each individual truth neuron and corresponding output neuron. **(b)** Same as a for a real recording (dataset d553101). Here the truth is only known for one cell thus only the performance relative to this cell can be computed.

### Performance of the network on simulated and real extracellular data

Figure 4 shows two examples of spike sorting results obtained on both artificial data with known ground truth and real extracellular tetrode recordings associated with an intracellular recording providing ground truth for one cell^32,33^ (see methods). Different types of error can occur when comparing the grown truth spike trains to the spike trains predicted by the sorting method: false negative, false positive and wrong classification. The performance of the method was assessed using the F-score (see methods), which accounts for all types of error. Figure 4a shows and example of 10 seconds of simulated data embedding 3 different action potential waveforms, firing at known time stamps according to a Poisson distribution with an average rate of 3.3 Hz. As shown in this figure, the output spiking pattern of the STDP network showed 3 mainly active output neurons whose spike trains closely resembled those of the three embedded signal waveforms. The network quickly converged to provide high classification rates as assessed over the last 100 s by an overall F-score of 85%, with individual scores of 90, 89, and 93% for each of the waveforms. Figure 4b shows an example of 10 s of real extracellular data for which the activity of one neuron was known from a simultaneous intracellular recording. The output of the STDP network showed 3 main active neurons, the activity of one of which was close to that of the intracellularly identified cell. For this cell, the sorting F-score of the network was stable for 150 s with an average value of 89% on the last 100 s.

We further compared our method with two open-source software able to perform unsupervised spike-sorting: Osort^28^ and Wave_Clus^22^. Figure 5a shows confusion results for the three methods on a simulated dataset. In this example, the STDP network made only a few false negative errors, while Wave_Clus made both false negative and false positive errors and Osort failed to detect one of the waveform and misclassified the two others in the same cluster, with an important number of false negative errors (Figure 5a). We further considered artificial data with varying signal-to-noise ratios (SNR). At very high SNRs, the STDP method gave slightly lower performance than Osort and Wave_Clus, but the F-score differences remained low (<0.06). At lower SNRs more comparable to those encountered in real cortical recordings, the STDP method became more efficient than the two other methods with strong F-score differences up to 0.3 (Figure 5b). This result was further observed on real neural signals (Figure 5c). In this case, the STDP approach performed similarly or better than Wave_Clus in all cases and performed better than Osort at low SNR.

**Figure 5.**
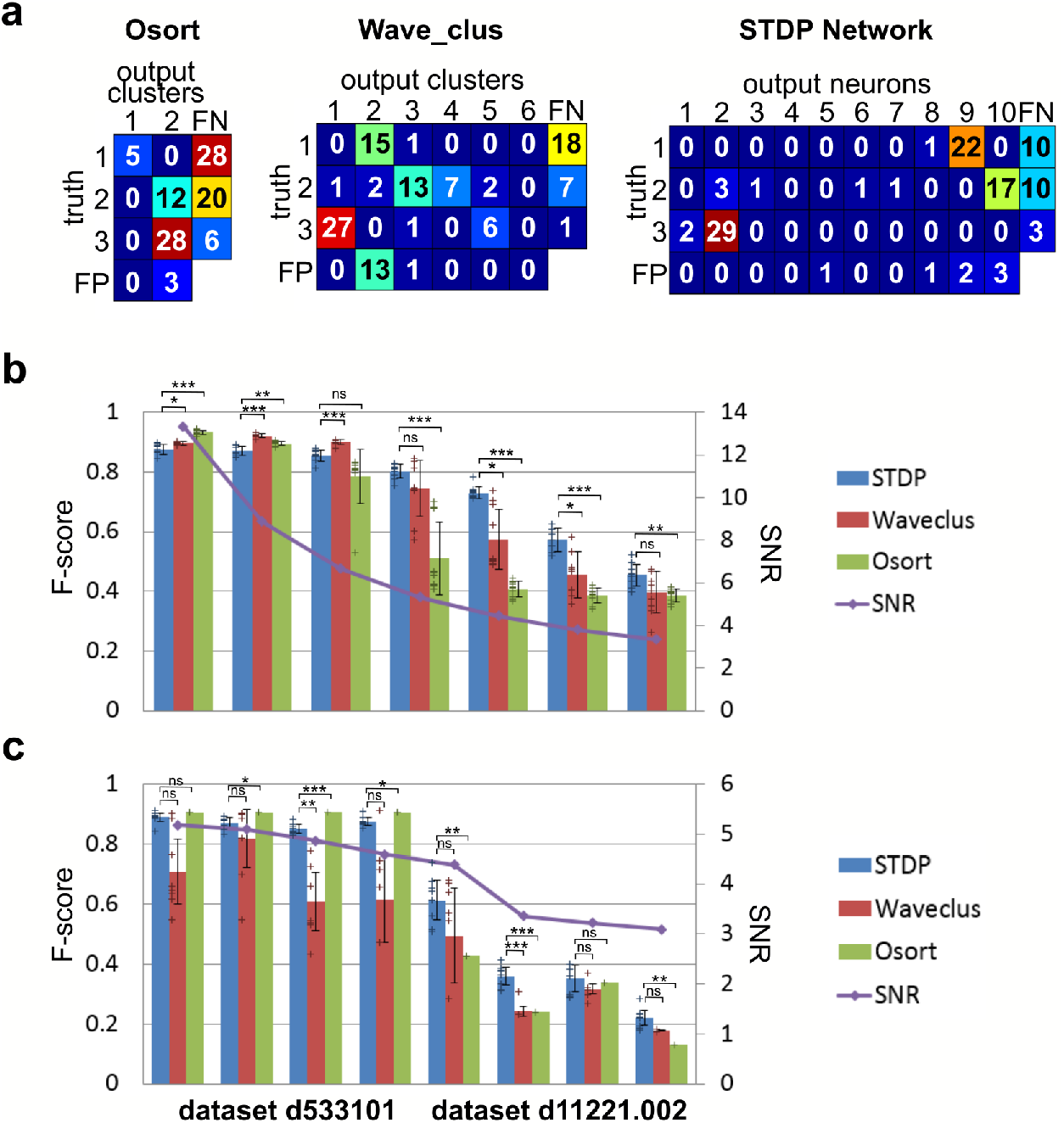
Comparison of the STDP network to two other algorithms (Osort and Wave_Clus). **(a)** Confusion matrices for the 3 sorting methods for a SNR=4.4 simulated dataset. The values are regularized relatively to 100 ground truth spikes i.e. 33 occurrences of each waveforms. **(b)** Mean F-score obtained on simulated data. Different SNR are considered, and for each SNR the performance was averaged over 10 different datasets, containing the same three waveforms. **(c)** Mean F-score obtained on real tetrode recordings sorted by decreasing SNR. For each channel, the STDP network was run 8 times, and Wave_Clus was run 8 times. The performance was averaged over the different runs. As Osort gives deterministic results, it was run only once for each channel. In b and c, the bars show the mean performance, the error bars show the standard deviation and the crosses show all data points. The purple curve shows the SNR of the tested datasets. The statistical test used for comparison was a Welch test, except for the comparison with Osort on the real data, where a one-sample t-test was used. “ns” stands for non-significant, * for p<0.05, ** for p<0.01, and *** for p<0.001. A Bonferroni correction was used (14 comparisons for the simulated data and 16 comparisons for the real data).

### Stability of the network

A spike sorting method should generate robust classification irrespective of the level of activity of the cells. In particular, classification should be correct for highly active neurons as well as for poorly active neurons. We thus tested the STDP-based sorting method on artificial data where 3 neurons were simulated with different firing rates: 1Hz for one cell, 3Hz for another and 9Hz for the third cell (Figure 6a). The network successfully classified the three different neurons with an overall F1 score of 84%. Noticeably, the network learned faster the waveform of the most active cell and more slowly the waveform of the least active cell. We found that the network needed about 50 occurrences of a given waveform to reach a correct and stable classification. The proposed method is thus able to correctly classify different waveforms corresponding to neurons having different firing rates, as soon as each waveform has occurred enough times. A robust spike sorting method then requires that a classification remains correct when the firing rates of the cells fluctuate. Indeed neural recordings during behavioral tasks induce such fluctuations depending on the involvement of the cells into the tasks. We thus also tested the STDP-based sorting method on artificial data where 3 neurons had intermittent firing activity (Figure 6b). All cells had a firing rate of 3 Hz but one became active only 50 s after the others and then these two other cells became silent one after the other before all became active again. We found that these firing rate variations did not impair the algorithm performance (Figure 6b). In particular, after the network had stabilized learning of two waveforms, it could still learn a new one. Moreover, when a waveform that the network had learnt became transiently absent in the signal, it could still classify it immediately and successfully as soon as it reappeared, meaning that the network keeps memory of the waveforms that have been learned.

**Figure 6.**
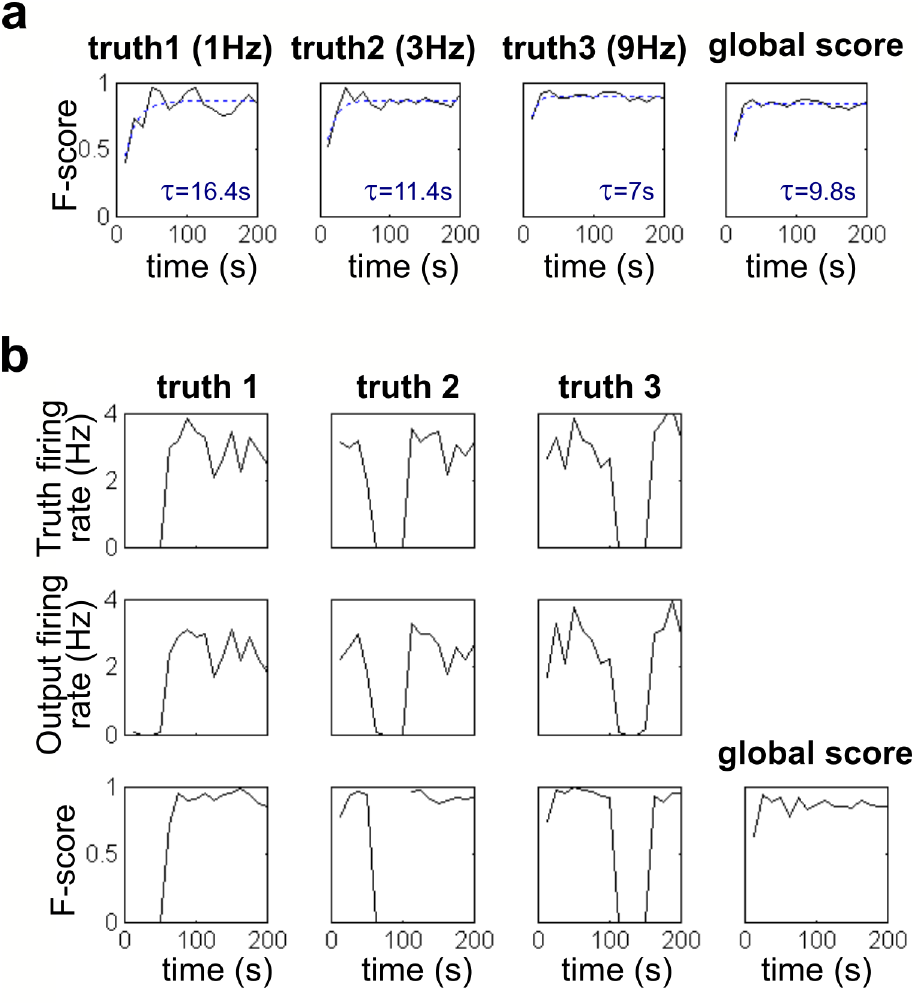
Robustness of the network to different firing rates scenarios. **(a)** Performance of the network along time for ground truth neurons with different firing rates. An exponential fit is performed to evaluate the duration needed for the network to learn each waveform. The network learns first the most frequent waveform 3, then waveform 2 having an intermediate firing rate and lastly the quietest waveform 1. **(b)** Evolution of the performance of the network for neurons with varying firing rates. Top: firing rate evolution of the 3 ground truth neurons: neuron 1 is initially silent, while neurons 2 and 3 get silent transiently one after the other. Middle: firing rate of the corresponding output neurons. Bottom: performance the network along time, for each ground truth neuron and for the overall population. The network performance remains stable despite the varying firing rates.

### Extension of the network to tetrode recordings

The recent advances of neural interfacing has benefited from novel micro and nanotechnologies allowing the fabrication of high-density multi-electrode systems^16,34–36^. Dense arrays of microelectrodes may provide neural recordings displaying partially overlapping information from one electrode site to the next. In particular, when recording sites are separated by less a few tens of microns, action potentials from a same cell can be recorded on several sites and spike sorting methods can benefit from combining neighboring electrodes instead of processing recording sites independently^23,27,37,38^. In this context, we thus adapted our method to the case of tetrode recordings, processing all channels at once instead of separately. To do so, we considered two variations of our network architecture. The first one had four input layers processing in parallel each of the four input signals and projecting to one common attention neuron and one common intermediate layer (Figure 7a). The second configuration had four input layers, four attention neurons and four intermediate layers working in parallel, the four intermediate layers and attention neurons projecting to one common output layer (Figure 7b). On the two datasets tested, we found that combining the four signals with the first configuration gave better classification than using each electrode separately. The second configuration, requiring a higher number of neurons and thus more computations, was only better for the first dataset having a higher SNR (Figure 7c).

**Figure 7.**
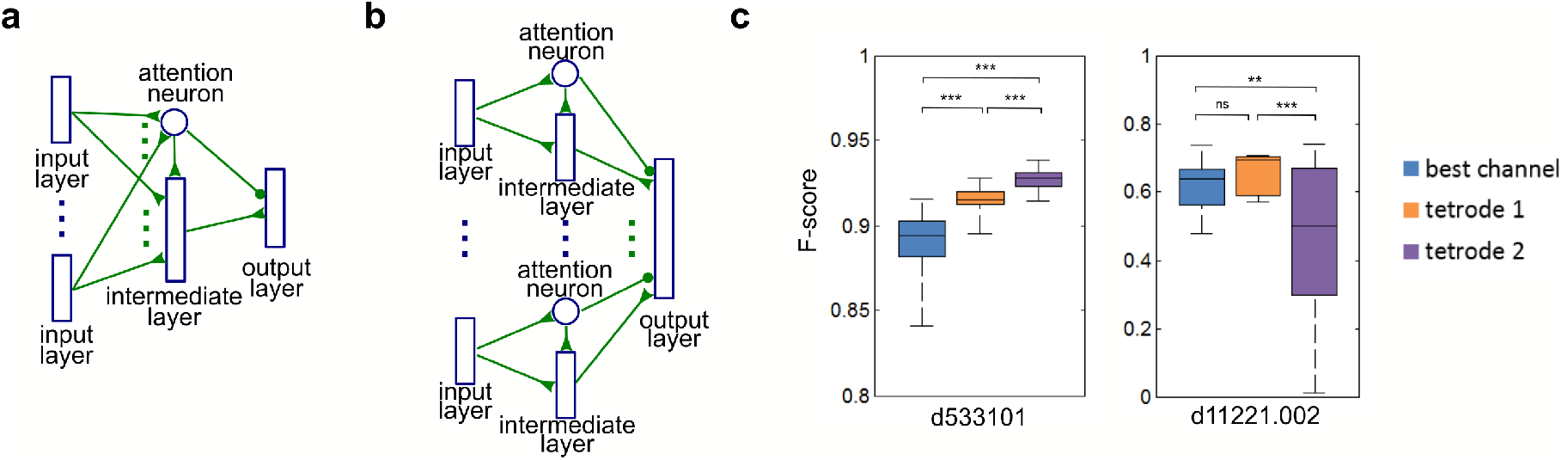
Adaptation of the network for the processing of tetrode data. **(a)** First adaptation (tetrode 1) where one input layer is used independently for each recording site and the intermediate and output layers are further shared. **(b)** Second adaptation (tetrode 2) where one input and one intermediate layers are used independently for each recording site and only the output layer is further shared. **(c)** Performance of the STDP network when using each channel separately and when using the 4 signals simultaneously with each of the two different architectures shown in a and b. The middle lines show the median performance, the box limits show the quartiles, and the whiskers show the extreme values. Each method was run 40 times on each dataset. The statistical test used for comparison was a Welch test. “ns” stands for non-significant, * for p<0.05, ** for p<0.01, and *** for p<0.001. A Bonferroni correction was used (6 comparisons).

## Discussion

Artificial STDP neural networks, inspired from biology^39,40^, are able to perform unsupervised learning^41^. Here, we considered such network to perform unsupervised spike-sorting. This choice was motivated by the development of neuromorphic circuits with memristive devices that offer a perspective to implement STDP networks on very low power embedded hardware^42^. In a preliminary study^43^ we showed that extracellular action potentials could be successfully sorted in case of a very high SNR. However, this method required an initial filtering of the signal by a bank of filters and became limited in case of realistic SNR. Here we propose a novel network architecture incorporating several plasticity rules showing performances comparable to existing methods with the advantage to process the continuous stream of data recorded by a microelectrode and directly outputting trains of artificial spikes representing the sorted activity of the recorded cells. It is further designed so that the input signal noise level was the only input-dependent feature influencing the parameters of the network.

In contrast to most spike-sorting algorithms that use separate steps for action potential detection, feature extraction and clustering, here the entire process is considered as a whole. The neural network directly processes the raw signal to output time stamps of sorted action potentials. This approach is thus directly compatible with online unsupervised spike sorting for real time processing of neural microelectrode recordings. Real-time processing however requires fast computations, especially when hundreds of recording sites need to be processed simultaneously. Here we demonstrated the method using conventional computer simulations without parallelization, and the computation time was definitely too high for a direct use in real time even for a single channel (computation time was here 5 to 10 times higher than the real time on a 3.2-GHz dual-core AMD Opteron processor). Yet, the method was designed to be compatible with future real-time embedding directly into large-scale microelectrode array systems. To this end, several implementation strategies can be considered. The algorithm could indeed be implemented with efficient parallelization on GPUs or dedicated Field-Programmable Gate Array (FPGA) hardware. Artificial STDP neural networks have indeed been implemented on FPGAs to demonstrate embedding neuromorphic computing^44,45^. However, although the real-time transposition of an artificial STDP spike-sorting network could be envisioned with GPUs or FPGAs, such strategies would still require high-power consumptions not compatible with future embedding in implantable devices. Future implantable devices embedding spike sorting at the electrode level will thus need to rely on other types of very low power implementations. The approach proposed here was designed specifically in such perspective. It indeed complies with neuromorphic circuits that currently undergo important developments. In particular scalable nonvolatile resistive memory (memristive) devices enable building artificial synapses with embedded STDP plasticity^29,42,46,47^ and thus open an opportunity to implement STDP networks in very power-efficient and miniaturized devices^43^ compatible with future implantable systems embedding neural signal processing.

The present work mainly covers the case of a single electrode signal (with an extension to the specific case of tetrodes). In order to process data recorded by a microelectrode array, this approach could be directly duplicated for each electrode independently if the pitch of the array remains large enough so that information is not redundant across recording sites. Yet, this is likely no longer pertinent for dense arrays offering continuums of electrodes with pitches typically less than a few tens of microns. In such devices, not only electrodes may ‘see’ several neurons, but neurons may also be seen by several electrodes. Here, we propose an adaptation of the method to the specific case of a tetrode, where the 4 recording sites can be combined altogether (Figure 7). However, when dealing with large dense arrays, there is no longer the notion of independent groups of electrodes: a group of electrode recording the action potential of the same neuron may well overlap with another group of electrodes recording another cell. Hence, specific methods should be developed where the array is considered as a whole^23,27^. Future work should thus focus on extending the proposed STDP network method to continuums of recording sites to comply with dense microelectrode arrays.

The present study may further open new ways to build hybrid neural networks. Such networks connect real neurons to artificial neurons. This field has been pioneered by dynamic clamp experiments at the single cell level using the patch-clamp technique^48^. Microelectrode arrays now open the way to build larger hybrid networks connecting multiple living cells to multiple artificial ones. Achieving a complete bi-directional hybridization requires two types of transformations: converting real neural network activity into artificial neural activity and, conversely, converting artificial activity into real activity. The artificial-to-real conversion has been achieved using electrical micro-stimulation at a network level^49^ but remains unachieved at the level of one-to-one artificial-real neuron pairs. Here, the spike-sorting STDP network offers a direct solution for the real-to-artificial one-to-one conversion by directly converting the real trains of spikes into a train of artificial spikes that become available to be further exploited by another downstream artificial neural network for further processing such as decoding algorithms. In particular, future rehabilitation approaches may benefit from combining such real-to-artificial hybridization with artificial-to-real microstimulation-based hybridization to achieve standalone fully embedded closed-loop devices.

## Acknowledgements

The authors wish to thank E. Vianello and J.-F. Bêche for helpful discussions.

## Online Methods

### Input layer

The input layer was organized into N_c_ columns corresponding to different processing delays of the input signal (see Figure 1b), regularly spaced by Δt_c_. Within each column, each neuron was sensitive to a given range of signal values. The size of this range, ΔV_s_, was the same for each neuron, and was set proportional to the noise level. Noticeably, the input signal noise level was the only input-dependent feature influencing the parameters of the network. The sensitivity ranges of neurons from one column are regularly overlapping so that, for each possible signal value, N_overlap_ neurons of this column fire. Within one column, the step between two consecutive sensitivity ranges was determined by the size of the sensitivity range ΔV_s_ and the overlapping factor N_overlap_, and the total number of neurons was determined by the total signal value range to cover. The input layer processed the input signal at regular time steps, Δts. This time step was chosen sufficiently short so that the time jitter due to sampling did not impact the performance of the network. As this time step was shorter than the original signal sampling time step, the signal was linearly interpolated for steps falling between samples. The values of the input layer parameters are given in Supplementary Table 1A.

### Attention neuron

The attention neuron was a Leaky-Integrate and Fire (LIF) neuron, which membrane potential V was governed by the following equation:

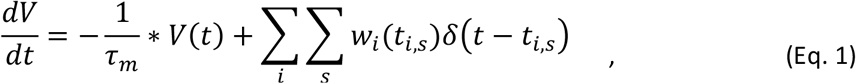

where V stands for the neuron potential, τ_m_ for the membrane time constant, i indexes the incoming synapses with synaptic weight w_i_, s indexes the received spikes from each synapse i with arrival time t_i,s_, and δ is the Dirac delta function. The membrane time constant of the attention neuron was chosen relatively short, in order to detect an action potential as early as possible. The neuron had no refractory period and no reset potential was applied. Instead, after firing, the potential was increased through a self-excitatory synapse, which constitutes a hysteresis mechanism (Supplementary Figure S2). The attention neuron threshold value was set empirically to obtain a compromise between false negative and false positive detection errors. The self-excitatory synapse ensured that, even if the action potential waveform crossed zero, thus lowering temporarily the attention neuron potential, the detection spike train was continuous from the beginning to the end of the action potential.

The weights of the synapses stemming from the input layer projecting to the attention neuron followed a short-term plasticity rule governed by the following equation:

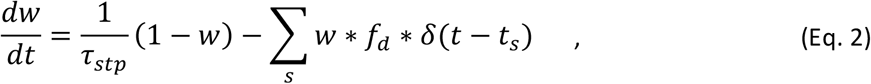

where w is the synaptic weight, τ_stp_ a time constant, f_d_ a depression factor, and t_s_ the presynaptic spike times. The time constant τ_stp_ was set long compared to the duration of an action potential, so that the synaptic weight did not change significantly during an action potential. The depression factor was set so that the asymptotic weight of a synapse stemming from an input neuron firing at each time step was significantly lower (x0.13 in our network implementation) than that stemming from an input neuron never firing.

The values of the attention neuron parameters are given in Supplementary Table 1B. For the processing of tetrode data where four input layers project to one attention neuron, the attention neuron threshold was multiplied by four.

### Intermediate layer

The intermediate layer was constituted of 60 LIF neurons following a similar equation as the attention neuron (Equation 1). We found that 60 neurons were enough to learn the different possible input patterns, which was confirmed by the fact that some intermediate neurons did not learn any pattern. It is possible to increase this number if a great variety of waveforms is expected in the signal. After firing, an intermediate neuron could not fire again during a refractory period τ_refrac_. After each firing or each lateral WTA inhibition, the neuron’s potential was reset to zero. The weight of the synapse coming from the attention neuron and the threshold of the intermediate neurons were chosen so that the intermediate neurons’ potential could not reach the threshold if the attention neuron was not firing, both before learning and after learning, and so that the time between two intermediate layer spikes was approximately Δt_c_. The feedforward synaptic weights could vary between 0 and 1 and were initialized randomly between 0.4 and 1 according to a uniform distribution. The initial average weight was thus 0.7, which is just enough to reach the threshold when the attention neuron is firing.

The STDP rule was based on a coincidence time window defined by two values, τ_stdp_+ and τ_stdp-_. A presynaptic spike was considered to coincide with a postsynaptic spike if it arrived at most τ_stdp_+ before the postsynaptic spike or at most τ_stdp-_ after the postsynaptic spike (see Figure 2a). For synapses targeting intermediate neurons, τ_stdp-_ was set to zero. For each post synaptic spike occurrence, the synapse weight was decreased by Δw_post_, and additionally, if a presynaptic spike coincided with the postsynaptic spike the synapse weight was increased by Δw_pair_ (resulting in a total change of Δw_post_+ Δw_pair_). The synaptic weight globally increased toward one if the ratio of pre- and postsynaptic spike coincidences over postsynaptic spike occurrences was superior to the ratio |Δw_post_/ Δw_pair_|, and decreased to zero otherwise.

The values of the intermediate layer parameters are given in Supplementary Table 1C. For the processing of tetrode data where four input layers project to one intermediate layer, the synaptic weight coming from the attention neuron and the intermediate neuron’s threshold were multiplied by four.

### Output layer

The output layer was constituted of 10 LIF neurons. The number of neurons was chosen higher than the maximum number of expected action potential waveform in the signal. These neurons had their potential reset after firing or after a lateral WTA inhibition. The reset potential and the lateral inhibition potential was proportional to the neuron threshold, which varied according to an intrinsic plasticity (IP) rule. Each pair of intermediate and output neurons were connected through multiple synapses with N_d_ different transmission delays and with one excitatory and one inhibitory synapse for each given delay. The attention neuron sent one inhibitory synapse to each synapse connecting an intermediate neuron to an output neuron. This inhibition was applied after the transmission delay (mimicking a biological presynaptic inhibition) so that intermediate spikes arriving at the output layer were prevented from being transmitted to the output neuron if the attention neuron was firing at the same time (see Supplementary Figure S3). These spikes were thus not taken into account for the STDP and IP rule.

The STDP rule used for the output layer (Figure 3a) was similar to the one used for the intermediate layer, with a different coincidence time window and different weight change values. The inhibitory and excitatory feedforward synapses followed an STDP rule with the same coincidence time window, but with a different |Δw_post_/ Δw_pair_| ratio, so that after learning, the summed weight converged to either 1, 0 or -1 depending on the ratio of the number of pre- and postsynaptic spike coincidences over the number of postsynaptic spike occurrences. Similarly to the intermediate layer, the initial weights were chosen randomly with a uniform distribution (see Supplementary Table 1D).

An important feature of the output layer was that the output neurons’ thresholds varied according to an intrinsic plasticity (IP) rule (Figure 3b). Every time an output neuron emitted a postsynaptic spike, its threshold Th was decreased by ΔTh_post_ = F^Δth^_post_*Th. For each presynaptic spike received within a coincidence window around the postsynaptic spike [-τ_IP+_ τ_IP-_], the threshold was increased by ΔTh_pair_ multiplied by the synaptic weight. The equilibrium threshold value was proportional with a factor F^Δth^_post_/ ΔTh_pair_ to the average weighted sum of the presynaptic spikes received within the coincidence window around a postsynaptic spike. The IP coincidence window was chosen according to the same proportion: τ_IP+_ / (τ_IP+_+τ_IP-_) = F^Δth^_post_/ ΔTh_pair_. The threshold was bounded between a minimum and a maximum values and was initialized at its maximum value (see Supplementary Table 1D).

The values of the intermediate layer parameters are given in Supplementary Table 1D. They remained unchanged when processing tetrode data.

### Simulated data

The spike sorting methods were first tested on simulated data generated using a method adapted from the literature^50^. Simulated signals were the superposition of correlated noise and action potentials occurring at known time stamps thus providing a known ground truth. The simulated sampling frequency was 20 kHz. The waveform of each truth neuron action potential was defined according to the following template equation:

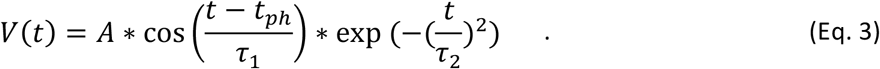

The parameters of this equation are given in Supplementary Table 2 for the 3 neurons that were simulated. The coefficient A was computed so that the maximum value of the template matched the parameter A_max_. The time occurrences of these waveforms were defined according to a Poisson process, with a mean firing rate of 3.3Hz each (unless otherwise stated), with a simulation time step of 0.05 ms corresponding to a 20 kHz sampling frequency. Additionally, a 3-ms refractory period was applied for each neuron. The noise added to the signal was a correlated noise generated by a dynamical Ornstein-Uhlenbeck process, with a 0.1-ms time constant and a simulation step of 0.002 ms. Simulated signal were generated with different noise levels to achieve different signal-to-noise ratios defined as 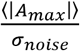, where 〈|*A_max_*|〉 was the mean of the peak amplitudes of the 3 action potentials and σ_noise_ was the standard deviation of the noise.

### Tetrode data

The spike sorting methods were also evaluated on real recordings from hippocampus region CA1 of anesthetized rats, available from the Buszaki Laboratory^32,33^ (datasets d533101 and d11221.002). Before sorting, the signals were band-pass-filtered using a first-order butterworth filter (300Hz – 3000Hz). The d533101 dataset, having an original 10-kHz sampling frequency, was up-sampled at 20 kHz using a Wittaker-Shannon interpolation.

### Spike sorting performance evaluation

To assess the performance of the spike sorting method, we computed indices based on an F-score. For each pair of truth neuron i and output neuron j, we computed the F-score as:

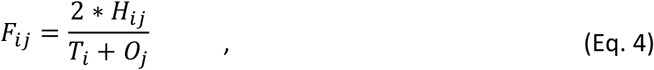

where T_i_ is the number of spikes emitted by the i^th^ truth neuron, Oj is the number of spikes emitted by the j^th^ output neuron, and H_ij_ is the number of output spikes coinciding with a truth spike within a 3-ms coincidence time window. Note that the F-score combines the recall and the precision of a prediction.

We also computed a global F-score across all truth neurons and all output neurons as:

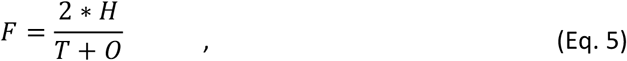

where T is the total number of truth spikes, O the total number of output spikes. To compute H, it is necessary to choose a correspondence between output neurons and truth neurons. Then H is the number of output spikes coinciding with a corresponding truth spike within a 3-ms coincidence time window. The correspondence between output neurons and truth neurons was chosen to maximize H.

### Statistical tests

The STDP network was compared to Wave-clus and Osort both on simulated data and on real tetrode data. For the simulated data, ten 200-s datasets were generated for each of the seven different noise levels, and each of the three compared methods were run once on each dataset. For the tetrode data, the STDP method was run 8 times on each channel, Wave-Clus was run 8 times on each channel, and Osort was run once on each channel as its result was deterministic. We then used the tetrode data to evaluate the performance of the STDP network when adapted to process all channels simultaneously. In this case, the STDP network was run 40 times both on each channel separately and on all channels simultaneously using two different tetrode methods (see Figure 7). Results obtained by the two tetrode methods were then compared to the best result obtained using only a single channel. As the variances were significantly different for the different groups (as assessed by a Bartlett test), a Welch test was used for 2-by-2 comparisons, except for the comparison with Osort on the tetrode data where a one-sample t-test was used. A Bonferroni correction was used for each set of multiple comparisons. All tests were performed using Matlab R2014a.

### Code availability

The custom code developed in this study is not made publicly available.

### Data availability

The artificial datasets generated and used in this study will be made available on Zenodo before publication. The real datasets used in this study are already available (see Ref 32).

---

**Supplementary Table 1:** implementation parameters

| A) Input layer parameters |  |  |
| --- | --- | --- |
| Parameter | Description | Value |
| $\Delta V_s$ | Sensitivity range size | $3.5\sigma_{\text{noise}}$ |
| $N_{\text{overlap}}$ | Number of neuron active at the same time within one column | 10 |
| $\Delta t_s$ | Time interval between two activations of the input layer | 0.0125 ms |
| $\Delta t_c$ | Time interval between two columns | 0.05 ms |
| $N_c$ | Number of column | 10 |
| B) Attention neuron parameters |  |  |
| Parameter | Description | Value |
| $\tau_m$ | Membrane time constant | $0.5 * \Delta t_s = 0.025$ ms |
| $\tau_{\text{refrac}}$ | Refractory period | 0 |
| $Th$ | Neurons threshold | $0.43 * N_{\text{overlap}} * N_c / (1 - \exp(-\Delta t_s / \tau_m)) = 108.1$ |
| $w_{\text{self}}$ | Weight of the self-excitatory synapse | $0.3 * (1 / \exp(-\Delta t_s / \tau_m) - 1) * Th = 10.5$ |
| $\tau_{\text{stp}}$ | Short term plasticity time constant | $10 * \Delta t_c * N_c = 5$ ms |
| $f_d$ | Short term plasticity depression factor | $1 - \exp(-6.5 * \Delta t_s / \tau_{\text{stp}}) = 0.0161$ |
| C) Intermediate layer parameters |  |  |
| Parameter | Description | Value |
| $N_{\text{neur}}$ | Number of neurons | 60 |
| $\tau_m$ | Membrane time constant | $2 * \Delta t_s = 0.025$ ms |
| $\tau_{\text{refrac}}$ | Refractory period | $\Delta t_c = 0.05$ ms |
| $V_{\text{reset}}$ | Reset potential | 0 |
| $V_{\text{inhib}}$ | Post-inhibition potential | 0 |
| $w_{AN}$ | Weight of the synapses coming from the attention neuron | $6 * \text{overlap} = 60$ |
| $Th$ | Neurons' threshold | $(w_{AN} + 7 * \text{overlap}) * (1 - \exp(-\Delta t_c / \tau_m)) / (1 - \exp(-\Delta t_s / \tau_m)) = 252.7$ |
| $w_{\text{init-}}$ | Minimal value for the feedforward synapses weight initialization. | 0.4 |
| $w_{\text{init+}}$ | Maximal value for the feedforward synapses weight initialization. | 1 |
| $\tau_{\text{stdp+}}$ | Positive STDP rule time window | $\Delta t_c + 0.5 * \Delta t_s = 0.0563$ ms |
| $\tau_{\text{stdp-}}$ | Negative STDP rule time window | 0 |
| $\Delta w_{\text{pair}}$ | Weight change for a presynaptic spike coinciding with a post synaptic spike | 0.005 |
| $\Delta w_{\text{post}}$ | Weight change for each postsynaptic spike | $-0.55 * \Delta w_{\text{pair}}$ |
| D) Output layer parameters |  |  |
| Parameter | Description | Value |
| $N_{\text{neur}}$ | Number of neurons | 10 |
| $\tau_m$ | Membrane time constant | 3 ms |
| $\tau_{\text{refrac}}$ | Refractory period | 3 ms |
| $F_{\text{reset}}$ | Reset potential factor. The reset potential is $V_{\text{reset}} = F_{\text{reset}} * Th$ | -10 |
| $F_{\text{inhib}}$ | Lateral inhibition potential factor. The lateral inhibition potential is $V_{\text{inhib}} = F_{\text{inhib}} * Th$ | -10 |
| $N_{\text{delays}}$ | Number of synaptic delays | 50 |
| $\Delta t_d$ | Time interval between synaptic delays | $\Delta t_c = 0.05$ ms |
| $w_{init-}^{pos}$ | Minimal value for the excitatory feedforward synapses weight initialization. | 0 |
| $w_{init+}^{pos}$ | Maximal value for the excitatory feedforward synapses weight initialization. | 1 |
| $w_{init-}^{neg}$ | Minimal value for the inhibitory feedforward synapses weight initialization. | 0 |
| $w_{init+}^{neg}$ | Maximal value for the inhibitory feedforward synapses weight initialization. | 0 |
| $\tau_{stdp+}$ | Positive STDP rule time window | 3 ms |
| $\tau_{stdp-}$ | Negative STDP rule time window | $4 * \Delta t_c = 0.2ms$ |
| $\Delta w_{pair}^{pos}$ | Excitatory synapses weight change for a presynaptic spike coinciding with a post synaptic spike | 0.05 |
| $\Delta w_{post}^{pos}$ | Excitatory synapses weight change for each postsynaptic spike | $-0.7 * \Delta w_{pair}^{pos}$ |
| $\Delta w_{pair}^{neg}$ | Inhibitory synapses weight change for a presynaptic spike coinciding with a post synaptic spike | -0.05 |
| $\Delta w_{post}^{neg}$ | Inhibitory synapses weight change for each postsynaptic spike | $-0.1 * \Delta w_{pair}^{neg}$ |
| $Th_{min}$ | Lower bound of the neuron threshold. | 8 |
| $Th_{max}$ | Upper bound of the neurons threshold, also used as initialization value | $0.35 * N_{delays} * Th_{min} = 140$ |
| $\tau_{ip+}$ | Positive IP rule time window | $6 * \Delta t_c = 0.3ms$ |
| $\tau_{ip-}$ | Negative IP rule time window | $4 * \Delta t_c = 0.2ms$ |
| $\Delta Th_{pair}$ | Threshold increase for each presynaptic spike coinciding with a postsynaptic spike | 0.01 |
| $F_{post}^{\Delta th}$ | Proportional threshold decrease for each postsynaptic spike. | $-0.6 * \Delta Th_{pair}$ |

**Supplementary Table 2:** Waveform template parameters

| | $A_{\max}$ | $\tau_1$ (ms) | $\tau_2$ (ms) | $t_{\text{ph}}$ (ms) |
| --- | --- | --- | --- | --- |
| **Neuron 1** | 5 | 1 | 0.5 | -0.25 |
| **Neuron 2** | 5 | 1 | 0.5 | 0.25 |
| **Neuron 3** | 10 | 1 | 0.5 | -0.19 |

**Supplementary figure S1.**
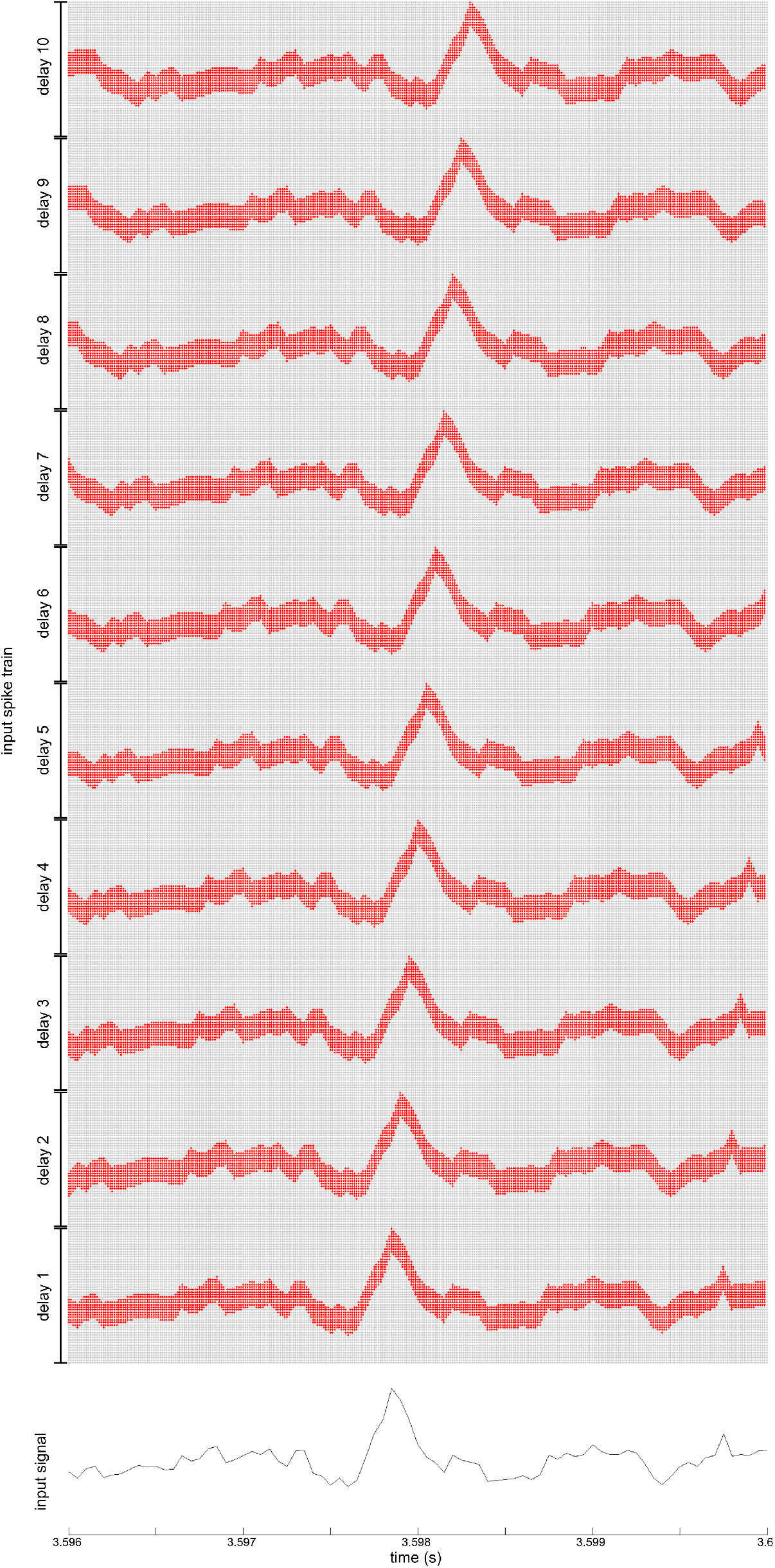
Encoding of the neural signal by the input layer. The lower curve shows an example of input signal. Above, the dot array represents the corresponding input layer activity. Each column of dots represents the activity of the input layer at one time step, the red dots are the active neurons and the gray dots are the silent neurons.

**Supplementary figure S2.**
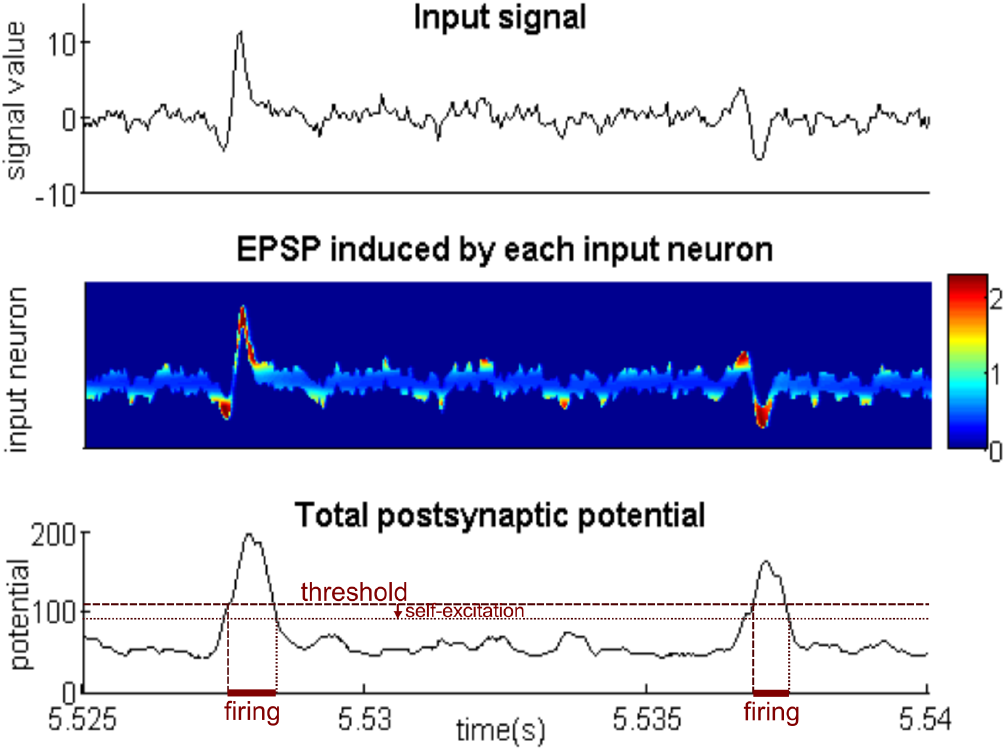
Short-term plasticity effect on the attention neuron. The upper panel shows an example of input signal. The middle panel shows the amplitude of the EPSP induced on the attention neuron by each input neuron of one input layer column. The frequently activated input neurons have a weak effect on the attention neuron whereas those rarely activated have a strong effect. The lower panel shows the time evolution of the attention neuron potential (when preventing it to fire by setting an infinite threshold): the potential of the attention neuron increases when an action potential occurs in the signal. Setting an appropriate threshold value (see Supplementary Table 1B) makes the attention neuron fire only when an action potential occurs. A hysteresis mechanism can be added by introducing a self-excitation.

**Supplementary figure S3.**
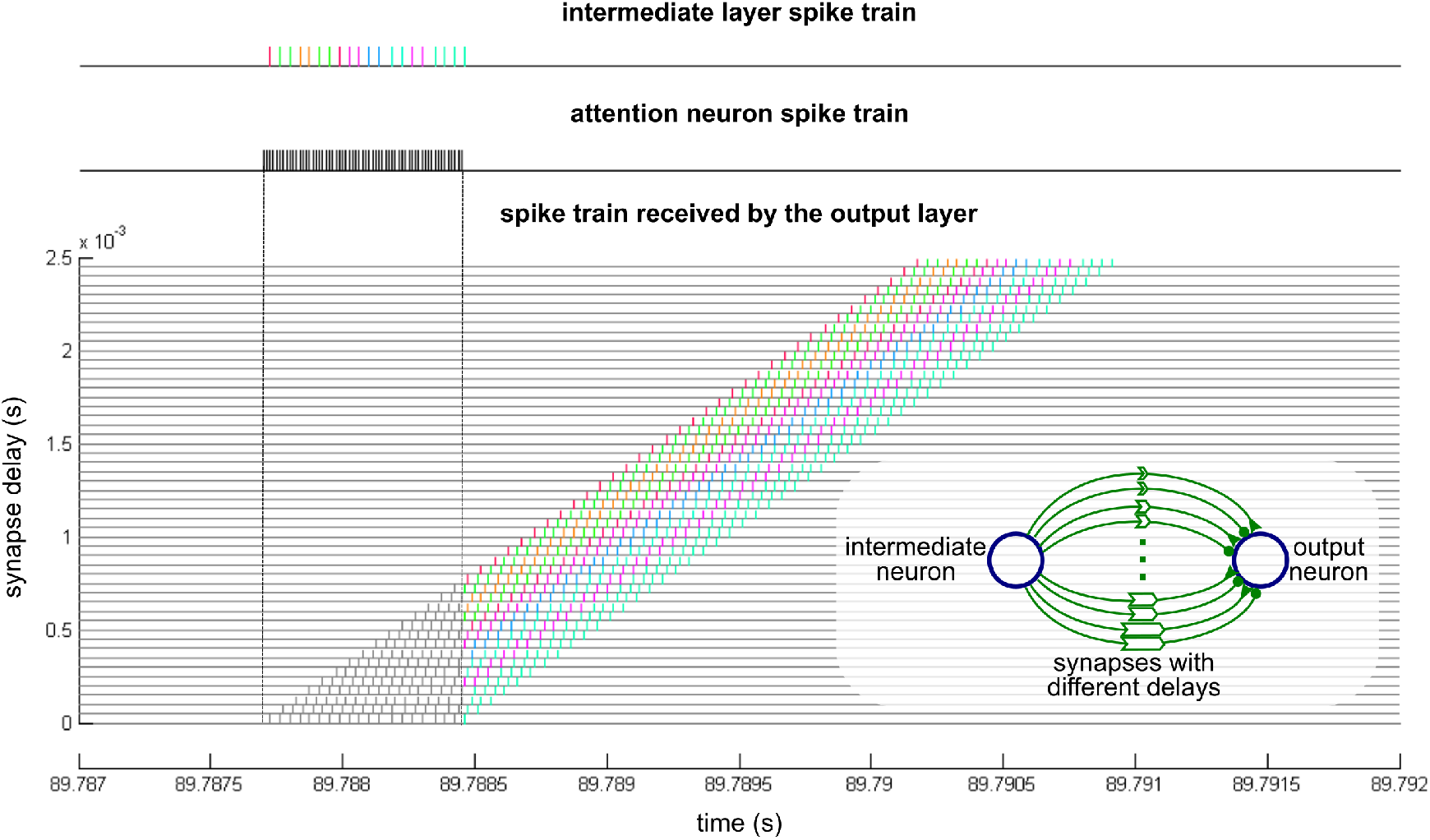
Synapses projecting from the intermediate layer to the output layer. The upper and middle traces show an example of attention neuron and intermediate layer spike train, generated by one action potential in the input signal (the different colors correspond to different neurons of the intermediate layer). The lower panel shows the spike train arriving on the output neurons, through synapses stemming from different intermediate neurons (in different colors) and with different transmission delays (vertical axis). As shown in the inset, each intermediate neuron is connected to each output neuron through several pairs of inhibitory (round endings) and excitatory (triangle endings) synapses having different transmission delays (represented by the length of the large arrows). Additionally the attention neuron prevents the intermediate spikes from arriving on the output neuron when it is firing, through a biological-like pre-synaptic inhibition on both excitatory and inhibitory synapse endings (see Figure 1a).

**Supplementary figure S4.**
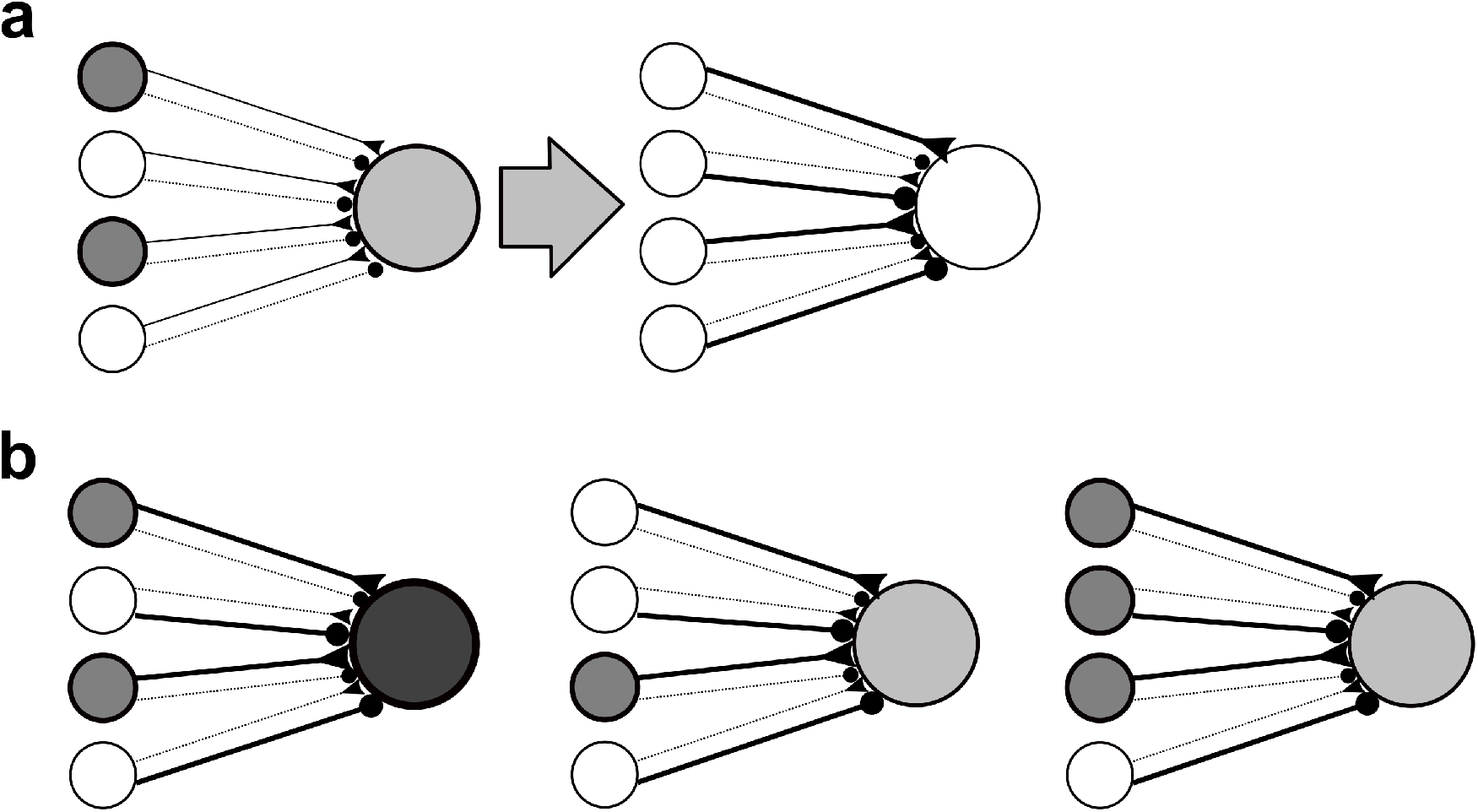
Role of inhibitory projections on the learning mechanism. Synapses with a triangular ending are excitatory, those with a round ending are inhibitory. Dotted lines represent weak synapses and bold lines strong synapses. The grey level of each neuron represents its excitation (white representing the lowest excitation) **(a)** When a postsynaptic neuron responds to a presynaptic pattern, the active excitatory synapses are potentiated while silent ones are depreciated. Conversely, the active inhibitory synapses are depreciated while silent ones are potentiated. **(b)** After learning the postsynaptic neuron potential is maximal when the learnt pattern is reproduced exactly (left). If the pattern is incomplete (middle), the output neuron is less excited. The output neuron is also less excited if an additional presynaptic neuron not belonging to the pattern is firing (right) due to its active inhibitory synapse.

